# Lessons Learned from a Ligand-Unbinding Stress Test for Weighted Ensemble Simulations

**DOI:** 10.1101/2025.04.23.650329

**Authors:** Anthony T. Bogetti, Darian T. Yang, Hannah E. Piston, David N. LeBard, Lillian T. Chong

**Affiliations:** Department of Chemistry, University of Pittsburgh, 331 Eberly Hall, Chevron Science Center, 219 Parkman Ave, Pittsburgh, Pennsylvania, 15260, United States of America; OpenEye, Cadence Molecular Sciences, Santa Fe, New Mexico, 87508, United States of America

## Abstract

The weighted ensemble (WE) path sampling strategy has pushed the boundaries of molecular simulation by enabling the generation of rates and atomistic pathways for biological processes beyond the ms timescale. However, the WE strategy has not yet reached its full potential and much can be gained from pursuing “stress tests.” Here we have explored a stress test involving the seconds-timescale unbinding of a highly charged ligand from a protein receptor: the release of the ADP ligand from the Eg5 protein receptor, which functions as a motor protein in cell division. From this stress test we learned valuable lessons regarding the choice of progress coordinate and improvements to the WE resampling procedure. Based on the latter, we have developed a WE method referred to as the Minimal Adaptive BinLess (MABL) method. The MABL method is in the same spirit as our previously developed Minimal Adaptive Binning (MAB) scheme for surmounting large energetic barriers, but is “binless”, i.e., does not require the use of rectilinear bins along a progress coordinate. This minimal version of a binless method is >50% more efficient than the corresponding binned version and provides a framework for implementing more complex binless methods.

## Introduction

The weighted ensemble (WE) path sampling strategy^1,2^ has enabled the generation of pathways with rigorous kinetics for grand-challenge applications in molecular simulation. These applications include large-scale conformational transitions within proteins^3,4^, protein folding^5–8^, protein-protein binding^9,10^, protein-ligand unbinding^11,12^, and chemical reactions^13^. The WE strategy efficiently samples barrier-crossing processes by running multiple, weighted trajectories in parallel and periodically applying a resampling procedure to provide even coverage of configurational space.^1,2^ Typically, configurational space is divided into regions, or “bins” along a progress coordinate—such as the distance between residues or the RMSD to a target structure—that captures the system’s slow, relevant motions. Although progress coordinates can be multi-dimensional, those with more than three dimensions are often impractical due to computational limitations.

To address this challenge, “binless” WE methods have been developed. These approaches reduce the cost of tracking progress in high-dimensional spaces by using a one-dimensional scoring functions instead of multi-dimensional progress coordinates.^7,8,14,15^ While binless WE methods still rely on metrics of progress, they apply these metrics in a more efficient way compared to binned WE techniques. A key feature of the WE strategy is that the progress coordinate can be changed “on the fly” during a simulation, as trajectory weights are independent of the specific progress coordinate used.^16^

Enhanced sampling strategies generally aim to make molecular simulations more efficient by reducing the time spent exploring redundant, stable states. These strategies promote broader sampling through one of three main approaches. The first class of strategies uses elevated temperatures to help systems overcome local barriers, with parallel tempering strategies—such as replica exchange molecular dynamics—being a widely used example. The second class introduces biasing forces to help the system surmount barriers, as exemplified by metadynamics. The third class, which includes WE and other path sampling strategies, enhances sampling of transitions between stable states rather than the states themselves without altering the free energy landscape or applying any biasing forces. Strategies in the first class cannot provide rate estimates. Methods of the second class—such as τ-random acceleration molecular dynamics (τ-RAMD)^17^, scaled-MD^18^, targeted MD^19^, and steered MD^20^—typically yield only relative rate estimates. Some approaches within the second and third classes, particularly metadynamics^21^, Gaussian accelerated MD^22^ and path sampling strategies more generally, can provide absolute rate estimates. The WE strategy is unique among these methods in its ability to provide absolute rate estimates while remaining highly flexible.^11^ Unlike post-simulation methods such as Markov state modeling or path sampling strategies such as weighted ensemble milestoning (WEM) that prioritize rate estimates with discontinuous trajectory segments,^23,24^ the WE strategy emphasizes the generation of continuous pathways. These detailed pathways not only yield rate estimates but also enable rich mechanistic insights.

Like any enhanced sampling method, there is no “free lunch” for WE path sampling over the conventional “brute force” manner of running sufficiently long molecular dynamics (MD) simulations to capture the process of interest. The main caveat of the WE strategy is that key motions of the process of interest may be orthogonal to the progress coordinate and sampled in a brute-force manner. That said, effective progress coordinates have been identified for a variety of complex biological processes on the seconds-timescale or beyond. For example, WE simulations have generated pathways and rates for a process involving the escape of an uncharged drug-like ligand from a completely buried cavity of a protein receptor.^11^ These simulations involved a combination of progress coordinates monitoring (i) the opening of the receptor cavity using the solvent accessible surface area of the cavity, (ii) the relative orientation of the ligand and receptor using an “unbinding” root mean squared deviation (RMSD) involving the heavy-atom RMSD of the ligand after alignment on the receptor in the bound state, and (iii) the ligand-receptor distance using the minimum distance between the two binding partners. To efficiently surmount barriers, bins were adaptively positioned along these coordinates using the minimal adaptive binning (MAB) method.^25^

Here we present lessons learned from a ligand-unbinding application in which the above WE protocol for uncharged ligands failed to generate successful events for a charged ligand. Our application was a “stress test” for the WE strategy and involved the unbinding of a highly charged ligand from a protein receptor, i.e., the release of an ADP ligand from the Eg5 motor protein (**Figure 1**), which is the rate-limiting step in a key process of cell division.^28^ The generation of fully-atomic, continuous ligand unbinding pathways and the prediction of ligand k_off_ values has long been of interest to the drug discovery pipeline given that drug efficacy is often correlated with the inverse of the k_off_, i.e., residency time of a ligand in the receptor binding site.^29^

**Figure 1.**
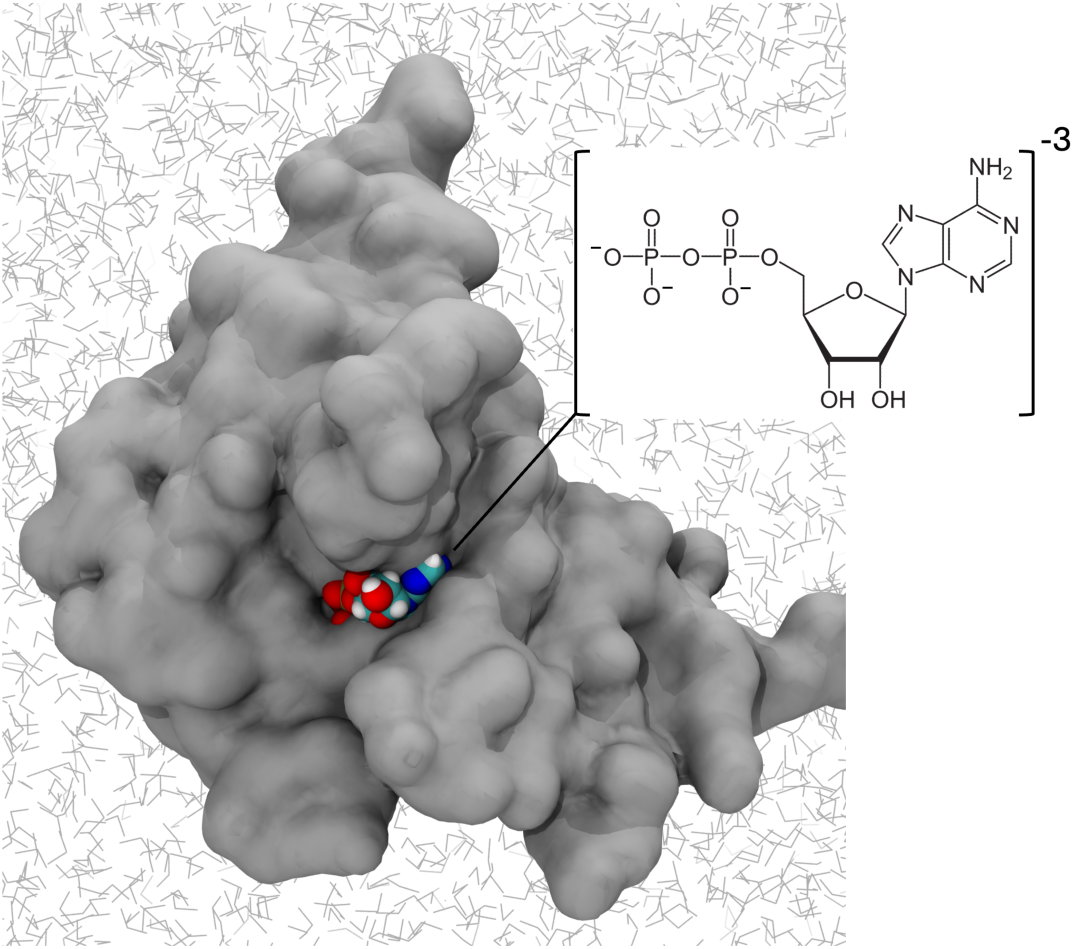
ADP-bound Eg5 motor protein. Crystal structure of the ADP-bound Eg5 motor protein (PDB code: 1II6).^26^ As a rate-limiting step for cell division, ADP release from Eg5 has been targeted for the design of allosteric inhibitors.^27^

The simulation of ADP unbinding serves as a relevant and challenging model system for developing methods to predict absolute k_off_ values. Our lessons learned provide insight into the choice of progress coordinate, manner of applying adaptive binning, and the WE resampling procedure. Based on the latter lesson, we have developed a minimal binless WE method (called MABL) that generalizes our MAB scheme.^25^ Our MABL method enables the efficient generation of continuous pathways for our highly charged ligand-unbinding stress test. The MABL method framework, along with many of the MABL variants tested for this project can be found at the following GitHub repository: https://github.com/atbogetti/MABL.

## Theory

In this section, we provide a brief overview of the WE strategy and outline the development of our minimal binless WE resampler, the MABL method, which allowed us to sample unbinding pathways for the highly charged ADP ligand. The MABL method is the extension of the MAB strategy for binless resampling, which provides greater flexibility and efficiency gains compared to the MAB method.

### The weighted ensemble (WE) strategy

The WE strategy involves running multiple weighted trajectories in parallel and periodic application of a resampling procedure at fixed short time intervals τ. This procedure involves replicating and terminating trajectories to provide even coverage of configurational space. Trajectories that populate less-visited regions of configurational space are replicated to enhance for success, dividing their statistical weight among the “split” trajectories. Trajectories that populate already-visited regions are terminated, merging their statistical weight with that of another trajectory in the same region. The combination of running MD for a resampling time interval τ and applying the resampling procedure constitutes a single WE iteration.

### The minimal adaptive binning (MAB) method with multiple regions

To efficiently surmount high energy barriers, we previously developed the minimal adaptive binning (MAB) method for adaptive placement of bins along a progress coordinate during a WE simulation.^25^ This method populates less-visited regions of state space by adaptively placing bins near the leading edge of progress and near bottleneck regions. Bins are also evenly spaced between the lagging and leading edges of the progress coordinate. Bin positions are determined after each resampling time interval τ.

For systems with particularly large barriers, one issue with the MAB method is the generation of trajectories with extremely low weights due to oversplitting—that is, repeatedly splitting the same leading trajectories. These very low weights can hinder convergence to a steady state, which is essential for obtaining reliable rate estimates. To reduce the likelihood of oversplitting, a multi-MAB scheme can be employed.^11,30^ In this approach, the progress coordinate is divided into several, relatively large regions and a separate MAB scheme is nested within each region (**Figure 2**). This use of multiple, independent MAB schemes is conceptually similar to other path sampling strategies, such as transition interface path sampling^31^ and the use of WE simulations between milestones in the weighted ensemble milestoning (WEM) method.^32^ The larger regions in which MAB bins are nested are defined by the user prior to the simulation and should reflect meaningful physical aspects of how the progress coordinates intersect. For example, in a recent WE application to the escape of an uncharged, drug-like ligand from a completely buried protein receptor^11^, the receptor cavity solvent-accessible surface area (SASA) needed to be binned more finely when unbinding RMSD was low, because at high unbinding RMSD the cavity would have had to already be open. Therefore, one MAB region was placed at low unbinding RMSD and another at high unbinding RMSD.

**Figure 2.**
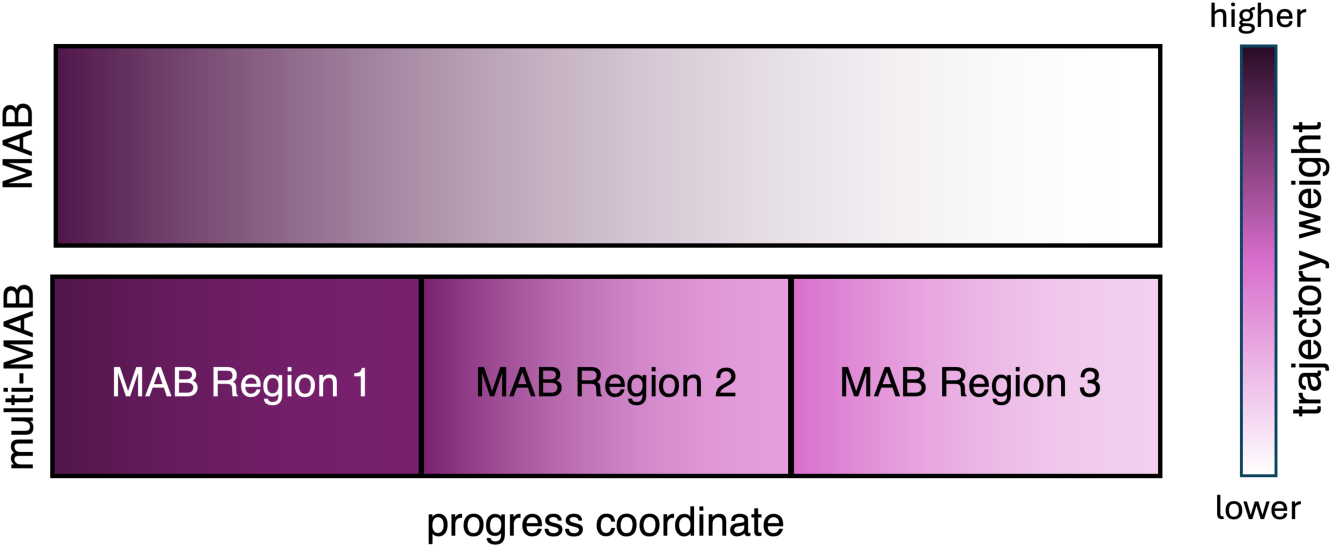
Basic illustration of the multi-MAB method. In the top schematic, a single MAB scheme (MAB method) covers the entire span of the progress coordinate, which can yield trajectories with extremely low weights due to oversplitting (i.e., repeated splitting of the same) leading trajectories. In the bottom scheme, the progress coordinate is divided into multiple regions, and a separate MAB scheme is nested in each region (the multi-MAB method). Nesting multiple MAB schemes in larger regions reduces the generation of leading trajectories with extremely low weights.

### The minimal adaptive binless (MABL) method

While the MAB method is efficient in surmounting large barriers, the computational expense of adaptive binning greatly increases with multi-dimensional progress coordinates. To enable efficient WE resampling of high-dimensional state space, we have developed a minimal adaptive binless (MABL) method for WE simulations. In contrast to binless WE methods such as the Resampling of Ensembles by Variance Optimization (REVO) method,^14^ which utilizes pairwise RMSD “distances” to split and merge trajectories, our MABL method is a minimal, intuitive binless scheme. Similar to our binned MAB method, the MABL method splits trajectories at the leading edge along one or multiple progress coordinates.

At the heart of the MABL method is the use of a progress score, denoted as S, which quantifies combined progress along multiple coordinates at the computational cost comparable to tracking a single coordinate. This score is calculated for each trajectory i after propagating dynamics for a short time interval τ, and reflects the trajectory’s advancement—on a scale of 0 to 1—along each m coordinate of progress q between an initial value q_m,i_ and target value q_m,t_ along M total coordinates:

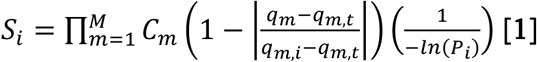

where q_m_ is the current value of coordinate m of trajectory i. The initial and target values of each coordinate are user-defined at the start of the simulation and can be adjusted on-the-fly during a simulation. The score S is calculated as a product of the progress along each individual coordinate, enabling the inclusion of multiple coordinates in a single metric. This allows for flexible and comprehensive monitoring of overall simulation progress. The scaling factors C_m_ enable users to tune the influence of each coordinate on the overall progress score. This adjustment serves a similar purpose to the multi-MAB scheme: to emphasize or de-emphasize certain progress measures depending on the expected relevance at difference stages of the simulated process.

To reduce the likelihood of oversplitting leading trajectories, a balance term 1/-ln(P) is included where P is the statistical weight of the trajectory being considered. By taking the fractional, negative log of P, lower-weight trajectories would be less favored for splitting, and higher-weight trajectories would be more favored for splitting thereby facilitating the flow of probability away from the initial state.

As illustrated in **Figure 3**, the WE resampling procedure in the MABL method begins by ranking all trajectories by their progress score S. To maintain a constant number of trajectories, the top N scoring trajectories are split, while the bottom N scoring trajectories are merged. Although the optimal value of N was not extensively benchmarked in our initial simulations, we found that N = 5 consistently facilitated the generation of productive ADP unbinding pathways. Based on this observation, we recommend choosing N to be ∼10-20% of the total number of trajectories in the WE simulation. It is important to note that N cannot exceed 50% of the total number of trajectories, as a single trajectory cannot be selected for both splitting and merging in the same resampling step. In accordance with WE resampling rules, when trajectories are merged, the surviving trajectory is selected probabilistically by the statistical weights of the candidates being considered.

**Figure 3.**
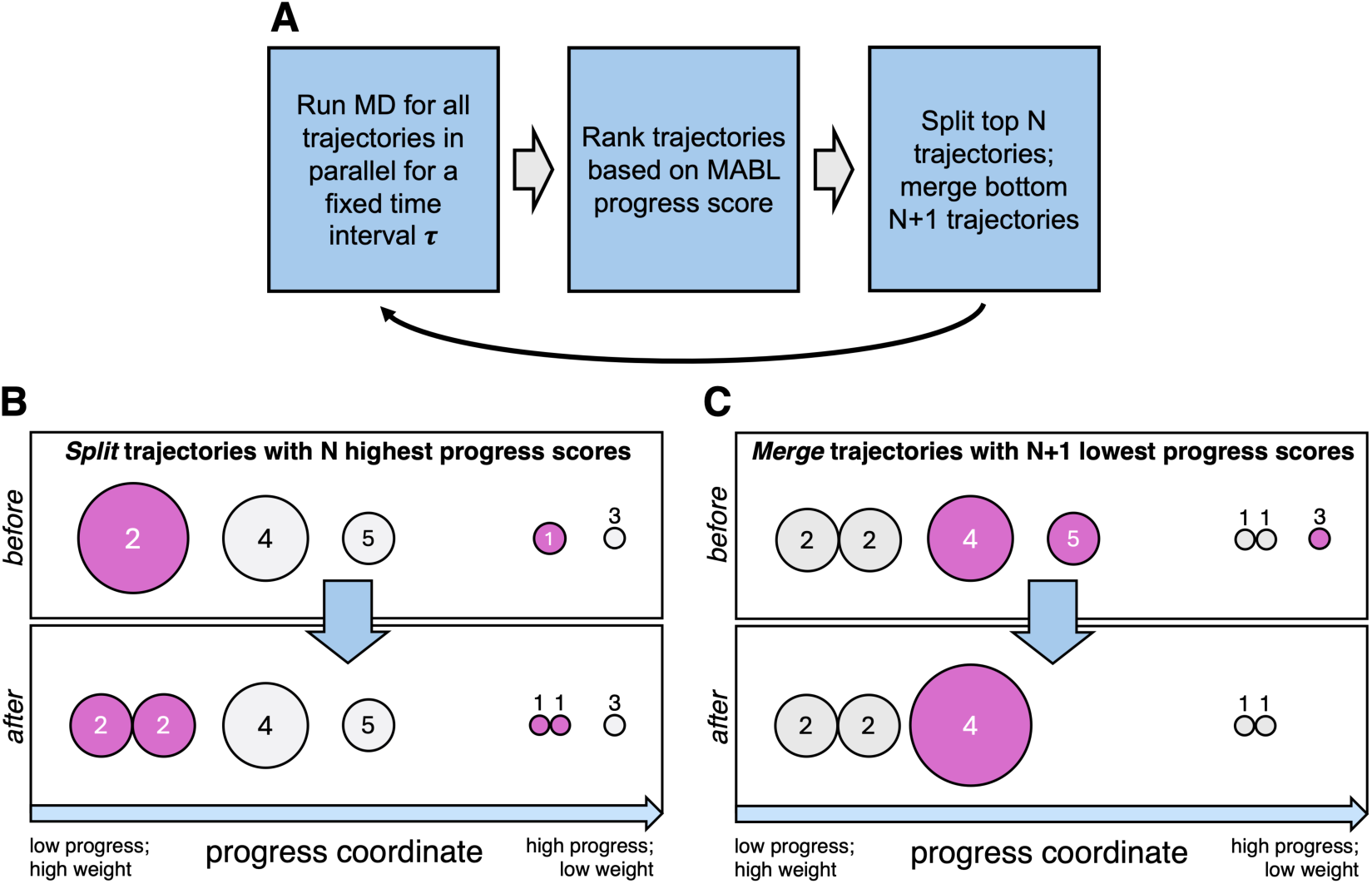
The MABL method for WE simulations. **A)** Workflow for the MABL method. **B)** Illustration of the splitting step in the WE resampling procedure. The ‘before’ box shows a trajectory distribution after tau, where trajectories with varying weights (represented by circle size) are ranked by a progress score S (see **Equation 1**; in this example, the only coordinate in S is the distance along the x axis). The top N = 2 trajectories (trajectories #1 and #2) are split, yielding the child trajectories in the ‘after’ box. **C)** Illustration of the merging step in the WE resampling procedure. To maintain a fixed total number of trajectories (here, 5 trajectories). the bottom N = 2 trajectories are merged.

Compared to a binned WE method, our binless MABL method greatly reduces the number of adjustable parameters for running a WE simulation, especially for more complex processes such as receptor-ligand unbinding. In particular, the use of a single progress score involves (i) defining only a range of values for the progress coordinate rather than dividing a progress coordinate into bins, and, optionally, (ii) specifying the relative importance of different progress coordinates rather than the complex task of nesting a progress coordinate within another progress coordinate.

## Methods

### Preparation of the initial state ensemble

All system preparation was performed using the tleap program of the Amber software package.^33^ The Eg5-ADP complex was prepared starting from heavy-atom coordinates of the complex from the crystal structure (PDB: 1II6)^26^, protonating titratable residues for neutral pH, and then solvating the system in a truncated octahedral box of explicit water molecules with a 16 Å clearance between the complex and the edge of the box. To mimic the experimental salt concentration^28^ of 100 mM NaCl, 73 sodium ions and 73 chloride ions were placed in the solvent box. The protein was treated with the Amber ff19SB force field^34^ and the waters with the OPC water model^35^. The ions, including a single Magnesium ion in complex with ADP, were treated with Li-Merz 12-6-4 parameters compatible with OPC water^36^ and the ADP ligand was treated with parameters compatible with the Amber ff19SB force field.^37^ Short-range interactions were truncated at 10 Å and long-range electrostatic interactions were treated using the particle mesh Ewald method^38^. The system was energy minimized for 10000 steps before heating for 20 ps with position restraints on all solute heavy atoms to 298 K at constant volume with a weak Langevin thermostat using a collision frequency of 1 ps^-1^. Following heating, the system was equilibrated for 1 ns with position restraints on all solute heavy atoms at constant pressure with a Monte Carlo barostat with pressure changes attempted every 0.2 ps. Finally, an additional 1 ns of constant-pressure equilibration was performed after removing solute heavy atom restraints.

### WE simulations

All WE simulations were run using the WESTPA 2.0 software package^39^, a resampling time interval of 50 ps, and initiated from the equilibrated bound structure of the Eg5/ADP system. All MD simulations were run using the pmemd.cuda GPU-accelerated dynamics engine of the Amber 22 package^33^ at constant temperature (298 K) and pressure (1 atm) using the weak Langevin thermostat and Monte Carlo barostat mentioned above for the equilibration procedure.

The multi-dimensional progress coordinate used in the MABL method incorporated the following metrics, each representing a different dimension of progress.

- **Unbinding RMSD.** The heavy-atom RMSD of the ligand, calculated after aligning the protein receptor to the initial bound-state structure. Alignment was restricted to receptor regions with relative low RMS fluctuations (residues 18-364), as identified by conventional MD simulations.
- **Ligand-receptor interaction energy (Eint).** The nonbonded interaction energy between the ligand and the receptor, including both van der Waals and electrostatic contributions. This was defined as the total energy of the ligand-receptor complex minus the sum of the individual energies of the ligand and receptor.
- **Ligand-receptor distance.** The minimum separation distance between any atom of the receptor and ligand.

This last metric was primarily used to define key states such as the unbound state, bound state, and if necessary, an encounter-complex intermediate. While it may have been conceptually cleaner to test each coordinate individually within the MABL framework, all three coordinates were ultimately necessary to capture the ADP unbinding process.

We compare our MABL WE simulation with a WE simulation run with a multi-MAB strategy. The multi-MAB WE simulation was run with a target number of 4 trajectories per bin. A four-dimensional progress coordinate was employed consisting of (i) the unbinding RMSD of the ADP ligand after alignment on the Eg5 receptor, (ii) the ligand-receptor interaction energy, (iii) the ligand-receptor separation distance, and (iv) the minimum separation distance between the phosphate tail of the ADP ligand and the Eg5 receptor. For each dimension of this progress coordinate, outer bin boundaries of [0, 7.5, 10.5, ‘inf’], [‘-inf’, 10 ‘inf’], [0, 6, ‘inf’] and [0, 6, ‘inf’] were defined.

Four MAB schemes were nested within the outer bins. The first MAB scheme was intended to sample increasing ligand RMSD and was placed at [3, 55, 5, 5] and contained [5, 1, 1, 1] MAB bins per dimension with directions of [1, −1, 1, 1]. The second MAB scheme was intended to sample increasing ligand-receptor distance to higher RMSD values and was placed at [8, 55, 5, 5] and contained [5, 1, 5, 1] MAB bins per dimension with directions of [1, −1, 1, 1]. The third MAB scheme was intended to sample interaction energies after RMSD had been increase over 10 Å and was placed at [11, 55, 5, 5] and contained [1, 5, 1, 1] MAB bins per dimension with directions of [1, −1, 1, 1]. The fourth and final MAB scheme was intended to focus sampling on increasing the minimum distance between the ligand’s phosphate tail and the receptor while also still focusing on sampling the interaction energy and was placed at [11, 5, 5, 5] and contained [1, 5, 1, 5] MAB bins per dimension with directions of [1, −1, 1, 1].

The MABL WE simulation was conducted with a fixed number of 40 trajectories per WE iteration. Progress scores in MABL were calculated using three quantities: 1) the RMSD of the ligand following alignment on the receptor, 2) the interaction energy between the ligand and receptor and 3) the minimum distance between the ligand and receptor. In contrast to the multi-MAB simulations, the minimum distance between the ligand’s phosphate tail and the receptor was excluded from the MABL simulations. This choice was based on extensive testing, which revealed that the metric was not necessary to achieve successful unbinding events. Since the metric was ultimately not needed, we do not expect it to have significantly impacted sampling efficiency in the multi-MAB simulations, and thus comparisons between MABL and multi-MAB simulations remain valid. For the RMSD, progress was evaluated between 0 and 25 Å; for the interaction energy, between 350 and −200 kcal/mol; and for the ligand-receptor distance, between 0 and 10 Å. In addition, to prevent the ligand from becoming “trapped” in nearby regions of the binding pocket, trajectories with RMSD values between 10 and 13 Å had their progress scaled by a factor of 0.8. These bounding values were selected based on preliminary WE simulations using a MAB scheme or multi-MAB scheme.

## Results and Discussion

In this section, we present in detail lessons learned from our successes and failures from WE simulations of a stress test: the unbinding process involving the charged ADP ligand from the Eg5 protein receptor. Among 44 different protocols for WE simulations of ADP unbinding from Eg5, we generated over 43.8 µs of aggregate simulation time. In this set of WE simulations, we tested a variety of progress coordinates, binning schemes and criteria for the WE resampling procedure. We learned two main lessons for applying WE simulations to study ligand unbinding that are particularly relevant to charged ligands. These lessons led to the development of our binless WE method, the MABL method, which was instrumental in enabling the efficient generation of ADP unbinding events.

<u>Lesson #1: Both energetic and structural features are necessary for monitoring unbinding of a charged ligand</u>

### Effective progress coordinates for the unbinding of uncharged ligands are not sufficient for charged ligands

While previous WE studies have found the unbinding RMSD to be an effective progress coordinate for simulating unbinding events with uncharged ligands,^11^ our tests revealed that this metric was not an effective progress coordinate for the unbinding of the charged ADP ligand. This result is due to the fact that the unbinding RMSD does not provide a continuous range of values that is sufficiently large to capture incremental amounts of progress toward the target unbound state. The detection of incremental amounts of progress are crucial for charged ligands given their tighter binding affinity for the protein receptor.

The WE strategy, when using a multi-dimensional progress coordinate, is most effective when one dimension has many “units” of progress towards the target state that simultaneously contribute to larger units of progress in other dimensions. For instance, in the case of receptor-ligand unbinding, increases in ligand-receptor interaction energy should (at least eventually) translate to increases in unbinding RMSD, which in turn translate to increases in receptor-ligand distance (the main, and most broad, determinant of success). In cases where the ligand is uncharged, unbinding RMSD may be sufficient as a progress coordinate because progress in unbinding RMSD can directly contribute to progress in receptor-ligand distance. However, in the case of charged-ligand unbinding, the tighter interactions between ligand and receptor compress the range of progress “units” available to the unbinding RMSD, rendering it not as effective. To overcome this, we identified and employed a more relevant coordinate with an order of magnitude more progress units that the WE strategy could utilize: the interaction energy.

### Interaction energy is an effective progress coordinate for sampling charged-ligand unbinding

The ligand-receptor interaction energy, with a large range of possible values due to electrostatic interactions, was effective as a progress coordinate for charged-ligand unbinding. By considering the interaction energy along with the unbinding RMSD and the ligand-receptor distance, we were able to get close to generating ligand unbinding pathways with the MAB method but were heavily limited by the large computational cost of binning in multiple dimensions of the progress coordinate. With the inclusion of the interaction energy as a coordinate in the MABL progress score, we were finally able to generate ligand unbinding events (**Figure 4**).

**Figure 4.**
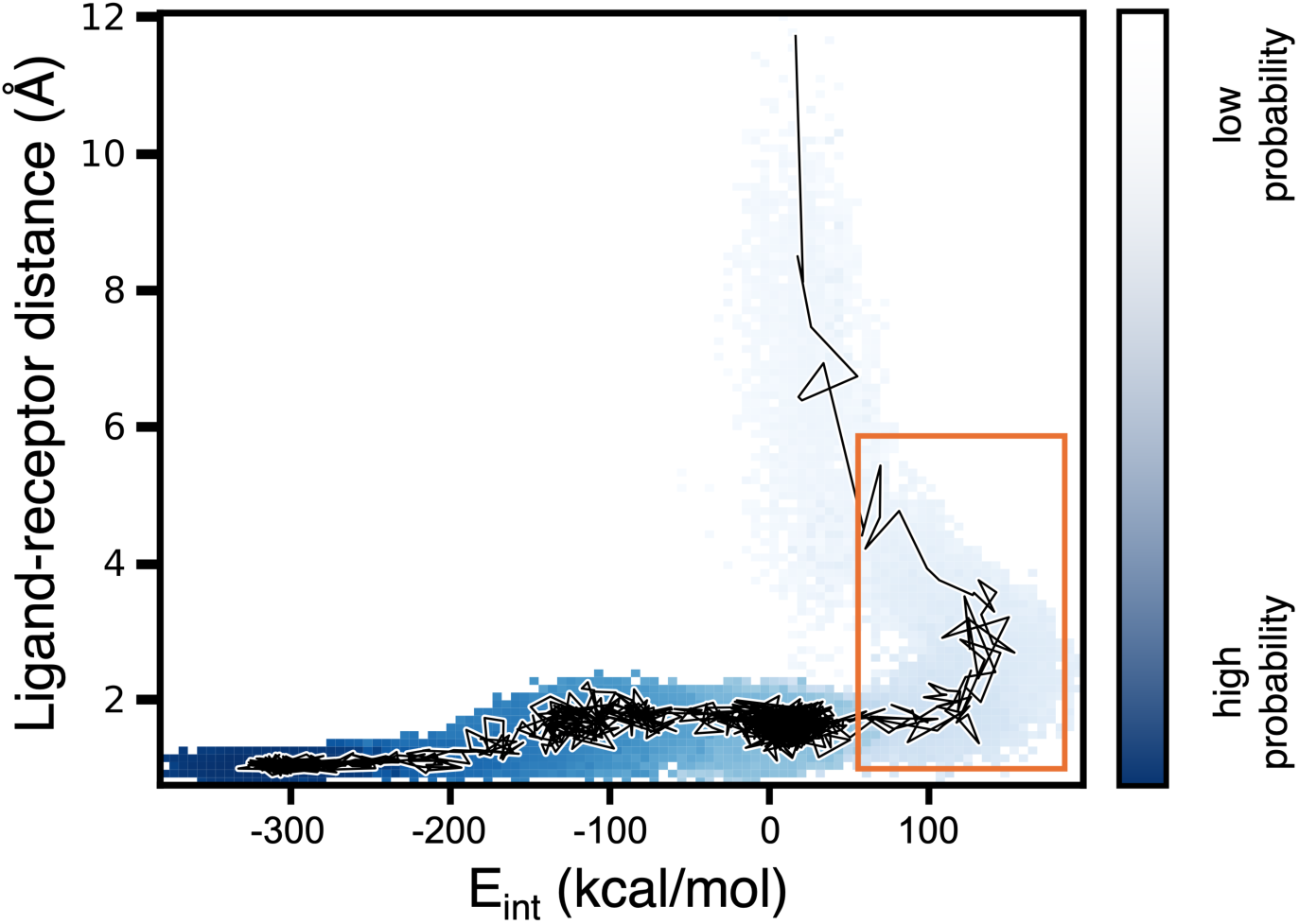
Repulsive ligand-receptor interactions facilitate ligand unbinding. Probability distribution as a function of the ligand-receptor interaction energy (E_int_) and ligand-receptor separation distance for a single WE simulation of the Eg5-ADP unbinding process. A representative unbinding pathway is traced in black. Repulsive ligand-receptor interactions (E_int_>0; delineated in red) appear to facilitate the dissociation of the ADP ligand from the Eg5 protein receptor. Given the extremely low probabilities in this simulation, which is far from a steady state, we interpret this distribution qualitatively.

### Backwards progress can be essential

As evident in **Figure 4**, repulsive interactions between the ligand and receptor occur immediately before the ligand dissociates from the receptor. Such interactions may serve as a “springboard” for launching the ligand away from the charged regions of the receptor binding pocket. Said another way, backwards motion along a progress coordinate can be not only useful, but essential for enhancing the sampling of pathways for unbinding of the charged ADP ligand.

In our initial efforts with the MABL method that employed a progress score consisting of the interaction energy, unbinding RMSD and ligand-receptor distance, our WE simulations resulted in trapping of the ligand-receptor system in conformations with large unbinding RMSD values. These trapped conformations corresponded to potential encounter complexes in which the ligand was bound to the receptor, but not in its native bound pose.

To enable successful unbinding events, we allowed backwards progress along the unbinding RMSD coordinate by reducing its relative contribution to the overall progress score by 20% in the range of 10 to 13 Å, where the ligand was getting trapped in a non-native bound conformation. After implementing this adjustment, the ADP ligand was able to fully dissociate from the Eg5 receptor, as shown in **Figure 4**.

<u>Lesson #2: While our binless WE method is more efficient in capturing rare events, care must be exercised in the merging of trajectories.</u>

### Our minimal adaptive binless (MABL) method enables efficient WE simulations with a high-dimensional progress coordinate

Using a progress score/coordinate that consists of the ligand-receptor interaction energy, unbinding RMSD, and the ligand-receptor distance, our binless MABL method was >50% more efficient at generating the first ligand-unbinding event compared to our most effective binned resampler employing the multi-MAB scheme. The aggregate simulation time needed for the MABL method to generate the first ligand-unbinding event was only 2.0 µs compared with 4.7 µs for the multi-MAB method. In addition, our MABL method maintains a fixed number of trajectories throughout the simulation—a desirable feature that facilitates the planning of computational resources to allocate for a WE simulation.

### Care must be taken when merging trajectories within a binless framework

While the progress score used in the MABL method is primarily focused on the splitting of promising trajectories, care must be exercised in the groupings of trajectories to consider for merging within a binless WE method for several reasons.

First, if non-redundant trajectories are grouped for merging, trajectories that are very different from the surviving trajectory could be terminated. The loss of non-redundant trajectories could hamper sampling of “breakout events” and generate a less-diverse path ensemble. This issue has been addressed for the original Huber and Kim WE framework in the recent development of an equal-weight resampler.^40^

Second, the total probability may become concentrated in just one or a few trajectories, leading to much lower-than-average probabilities of trajectories, resulting much lower-than-average probabilities for those at the leading edge of sampling. We were able to alleviate—but not entirely eliminate—this accumulation by incorporating trajectory weights into the progress score. Additional strategies to more effectively prevent large-scale probability accumulation will be explored in future work.

Based on our efforts to refine the WE resampling procedure in our MABL method, we present a cautionary example where we violated one of the two rules that must be followed by a resampling procedure to avoid introducing statistical bias into the WE simulation:

1. Trajectory weights must always sum to a total probability of one, and
2. When evaluating trajectories for merging, the surviving trajectory must be chosen according to its weight (probability) or randomly chosen, if more than one candidate exists with the same weight.^1^

In our example, we broke the second rule when attempting to select the surviving trajectory based on their progress score. While the progress score contains a weight-based balance term (see Methods), this criteria for selecting the surviving trajectory resulted in the termination of a higher-weight trajectory and the subsequent merging of its weight onto a lower-weight surviving trajectory. This merging event led to an unphysical result where there was no reduction in weights as the trajectories approached the target state (i.e., as if no barriers exist in the unbinding process) due to two high-weight trajectories ending up on a “fast track” from the initial to the target state (**Figure 5**). Our example underscores the importance of following not just the first rule, but also the second rule for the WE resampling procedure that involves the random selection of trajectories that survive a merging event.

**Figure 5.**
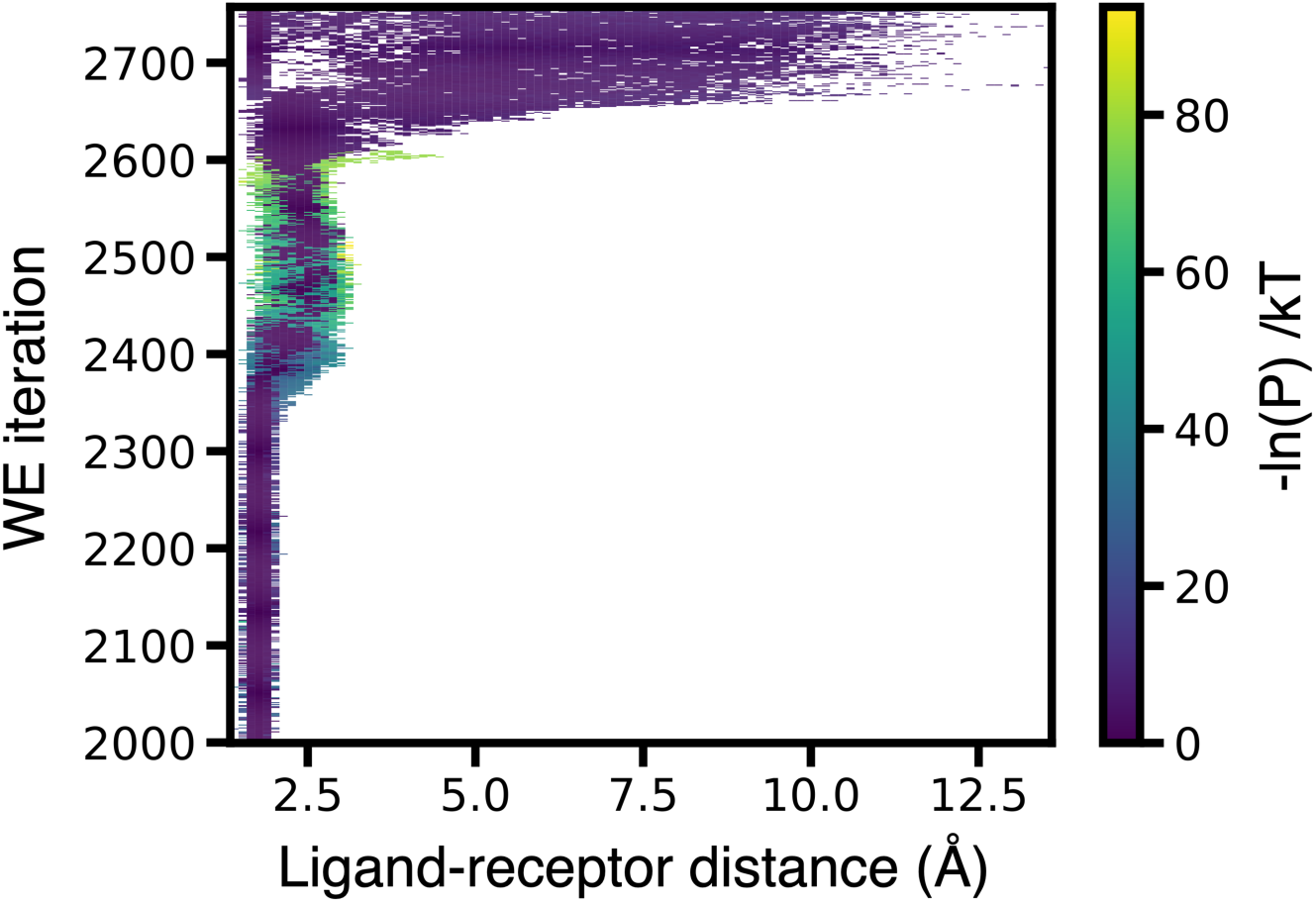
Time-evolution of the probability distribution as a function of ligand-receptor distance from a WE simulation in which the choice of trajectories for merging breaks the rules. The preferential merging of heavy-weight trajectories in an initial version of the MABL method violated the statistical rules of the WE resampling procedure.^1^ As a result, a few heavy-weight trajectories make a “bee-line” from the initial to target state without any splitting of the trajectories.

### A strategy for enforcing “soft” threshold values for trajectory weights

In certain WE studies, the use of minimum and maximum threshold values for trajectory weights has prevented the weights from becoming extremely small and dominating the path ensemble, respectively.^12,14^ However, hard limits on trajectory weights can hinder progress towards the target state by preventing any splitting of “breakaway” trajectories. To achieve a middle ground between being too strict and too relaxed with trajectory weights, we introduced “soft” thresholds into the MABL method by incorporating trajectory weights into the progress score. Based on this progress score, higher-weight trajectories are chosen for splitting and lower-weight trajectories are considered for merging. The “soft” threshold term used in our progress score can be further modified and serves as a starting point for exploring the use of this more flexible form of trajectory weight thresholds.

## Conclusions

We have presented lessons learned from WE simulations of a challenging stress test: the unbinding of a charged ligand, specifically the release of ADP from the Eg5 motor protein. Our lessons are drawn from >43.8 µs of aggregate WE simulation time across 44 different simulation protocols. Many of our unsuccessful attempts ultimately led to the development of a new WE strategy—the MABL method—a binless version of our previously developed MAB method.

The MABL method performs WE resampling using a single progress score that integrates progress along multiple coordinates. This binless approach proved more efficient than our best binned strategies when handling multiple progress coordinates. Further, MABL maintains a fixed number of total trajectories throughout a simulation, which simplifies computational resource allocation.

Despite the complexity of simulating the unbinding of a charged ligand, we successfully generated unbinding pathways using the MABL method by incorporating three key features: (i) inclusion of ligand-receptor interaction energy in the progress score, (ii) enabling backwards progress by reducing the weight of the unbinding RMSD coordinate where the ligand was prone to becoming trapped, and (iii) mitigating oversplitting at the leading edge of sampling by integrating trajectory weights into the progress score. Although rate constants could not yet be estimated from these initial pathways, they represent an important first step toward that goal.

Finally, we have implemented our MABL method within the open-source WESTPA software package, providing a framework for future development of more advanced binless strategies to address complex biomolecular processes.

## Author Contributions

The manuscript was written through contributions of all authors. All authors have given approval to the final version of the manuscript.

## ACKNOWLEDGMENT

We thank AJ Pratt for initial efforts and Anthony Nicholls for insightful discussions.

---

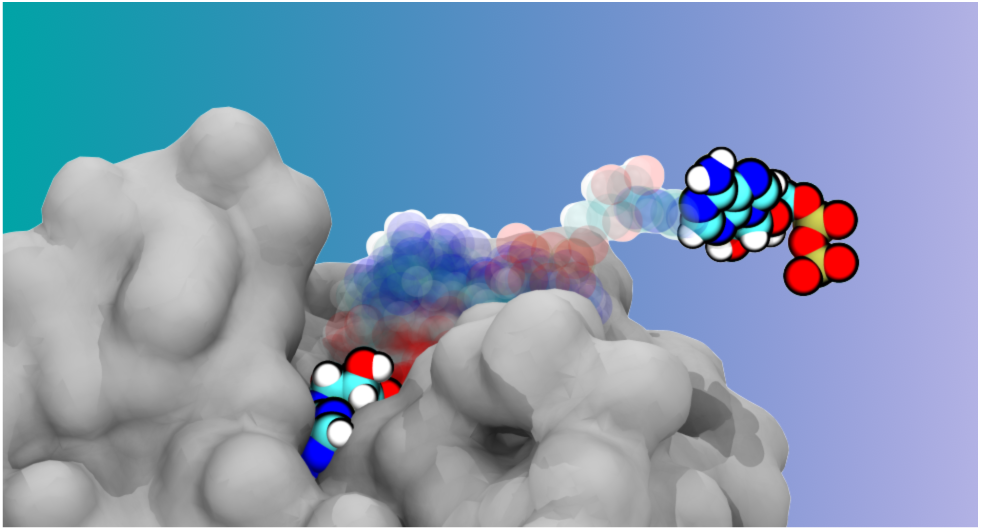
For Table of Contents Only.

